# SingleCellNet: a computational tool to classify single cell RNA-Seq data across platforms and across species

**DOI:** 10.1101/508085

**Authors:** Yuqi Tan, Patrick Cahan

## Abstract

Single cell RNA-Seq has emerged as a powerful tool in diverse applications, ranging from determining the cell-type composition of tissues to uncovering the regulators of developmental programs. A near-universal step in the analysis of single cell RNA-Seq data is to hypothesize the identity of each cell. Often, this is achieved by finding cells that express combinations of marker genes that had previously been implicated as being cell-type specific, an approach that is not quantitative and does not explicitly take advantage of other single cell RNA-Seq studies. Here, we describe our tool, SingleCellNet, which addresses these issues and enables the classification of query single cell RNA-Seq data in comparison to reference single cell RNA-Seq data. SingleCellNet compares favorably to other methods, and it is notably able to make sensitive and accurate classifications across platforms and species. We demonstrate how SingleCellNet can be used to classify previously undetermined cells, and how it can be used to assess the outcome of cell fate engineering experiments.

**Highlight:**

- SingleCellNet (SCN) enables the classification of scRNA-Seq data across platforms and species
- SCN is open source and extendible
- We illustrate the utility of SCN with three example applications

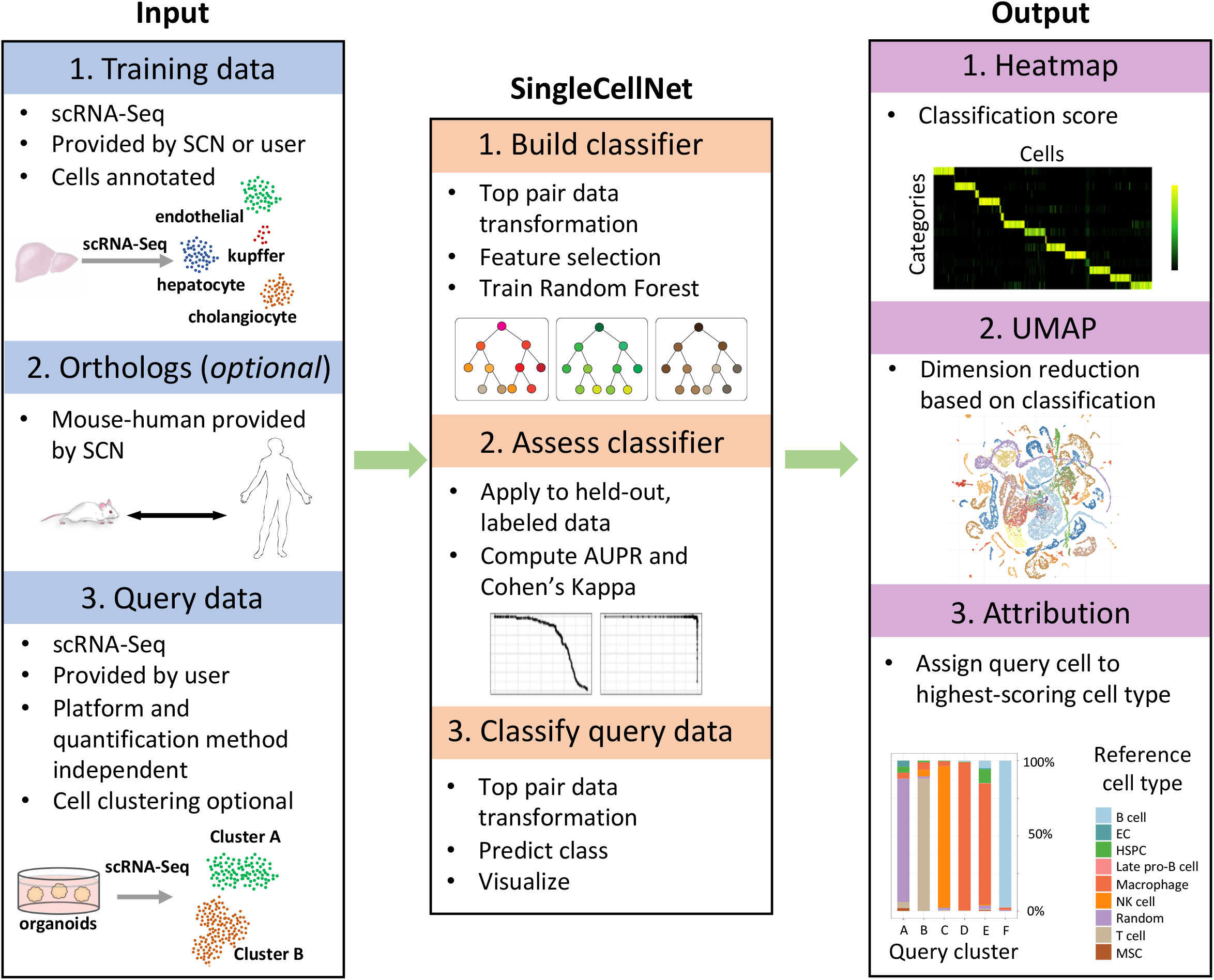

## Introduction

Single cell RNA-Seq (scRNA-Seq) has rapidly emerged as a powerful tool to generate cell atlases of organs, tissues, and even complete organisms (Cao et al., 2017; Han et al., 2018; Tabula Muris Consortium et al., 2018), to define stages and regulators of lineage commitment during development (Kumar et al., 2017), and to determine how perturbations such as age, pathology, or genetic variation impact cell composition and cell state (Haber et al., 2017; Kowalczyk et al., 2015; Park et al., 2018; Patel et al., 2014). One of the most time-consuming aspects of scRNA-Seq investigations is annotating each cell, or in other words, determining each cell’s ‘type’. This often requires further experimentation such as in situ-based methods to localize cells within a tissue, or prospective isolation followed by functional assessment. It is evident, that a faster method with more quantitative rigor method is needed. One approach would be to develop a method that allows the direct comparison of ‘query’ scRNA-Seq data to an existing scRNA-Seq dataset in which the cells have already been identified, such as a cell atlas.

Several methods to integrate scRNA-Seq data have been proposed. For example, canonical correlation analysis (CCA) (Butler et al., 2018), and MnnCorrect (Haghverdi et al., 2018) have proven useful in aggregating scRNA-Seq datasets so as to increase statistical power in differential gene expression analysis and in gene-to-gene correlation analysis. However, these approaches require that at least one relatively abundant cell type is present in both datasets. Furthermore, the methods do not explicitly provide a means to quantitatively classify query cell types in comparison to a reference data set, which is the goal of our method SingleCellNet (SCN). MetaNeighbor is another tool that compares cell types across scRNA-Seq datasets, yet it addresses the question “to what extent is a group of cells reproducible across scRNA-Seq data sets?”, which is distinct from our aim (Crow et al., 2018). SCMAP is the method most akin to SCN in intent (Kiselev et al., 2018) in that it classifies query cells according to their similarity to reference cell types based on various measures of correlation. While SCMAP is fast, it ultimately returns a binary cell type assignment for each cell. In many applications, such as in cell fate engineering, a quantitative measure of similarity would be more informative than a categorical assignment of identity. Here, we present SCN, a method to quantitatively classify scRNA-Seq data based on comparison to a reference data set. To make query and reference data compatible across platforms and species, we use a simple transformation based on comparing expression of pairs of genes within each cell, a method inspired by the top-scoring pair classifier (Geman et al., 2004). Here we evaluate the performance of SCN, compare it to the intermediate quantitative outputs of SCMAP, and highlight its utility in three realistic use-cases: a cross-platform identification of previously unclassified cells, the identification of cell types resulting from a cell fate engineering experiment, and a cross-species classification of hematopoietic cell types.

## RESULTS

### Building a multi-class scRNA-Seq classifier with top-pair transform and Random Forest

We previously developed CellNet, a computational method designed to classify populations of engineered cells (i.e. those derived by differentiating pluripotent stem cells) (Cahan et al., 2014) using Random Forests (Breiman, 2001). With SCN, we have revamped this approach to enable classification of scRNA-Seq data generated from different platforms and from different species. We do not use gene counts or expression estimates directly in training or in classifying. Rather, we transform the data into a binary matrix derived by pairwise comparisons of selected genes on a per cell basis, limited to genes that are preferentially expressed in each cell type defined in the training data, as well as those genes that are specifically under-expressed in each type (**Fig 1A**). Then, to limit the set of predictors for input to training the Random Forest classifier, we use template matching (Pavlidis and Noble, 2001) to find the most discriminating sets of gene pairs.

**Figure 1.**
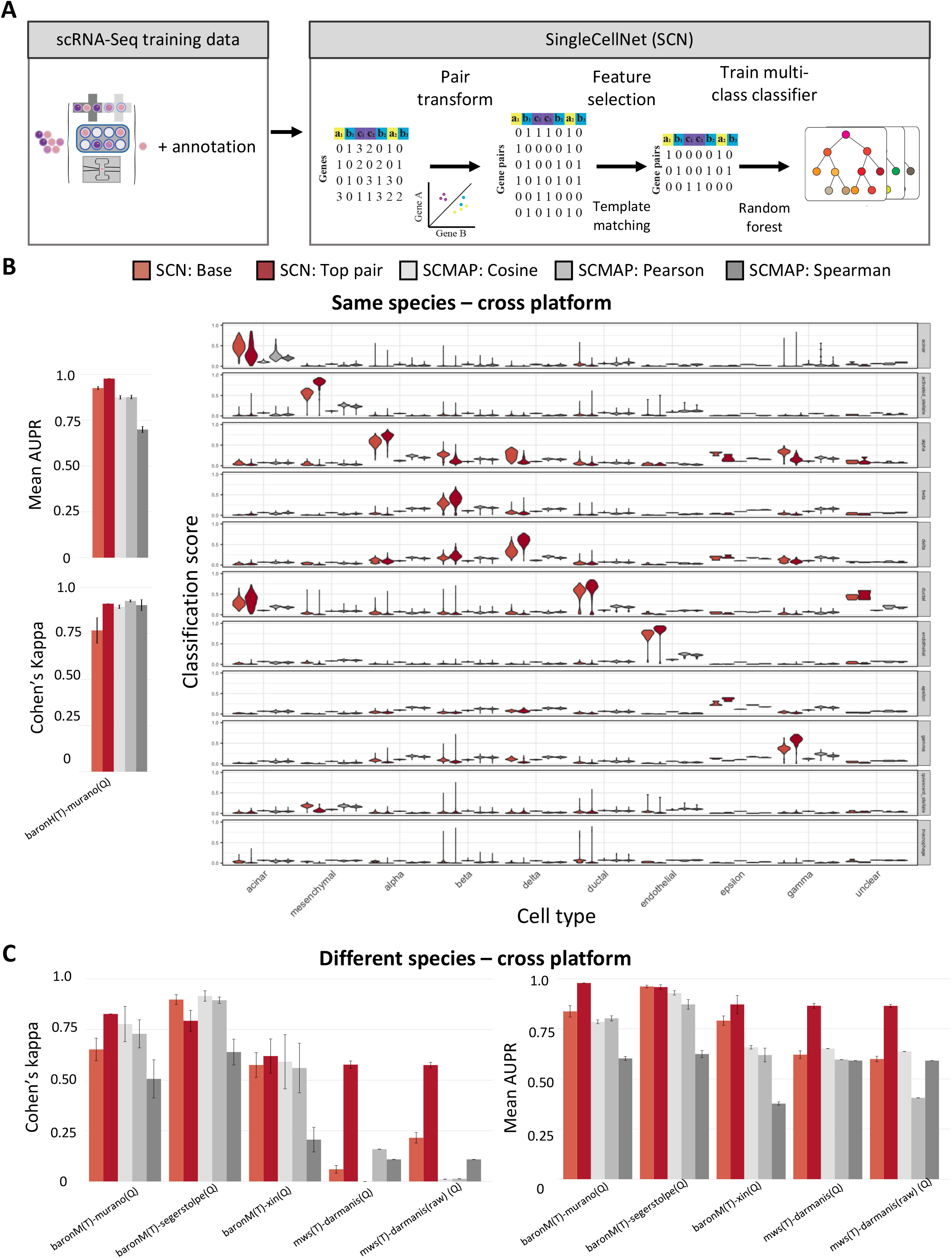
SingleCellNet (SCN) schematic and performance comparisons (A) SCN takes in scRNA-Seq with annotation as training data. SCN selects the best classifying gene pairs from the training data using an approach similar to the Top-scoring-pair algorithm and pair-transforms it into a binary matrix. A multi-class Random forest classifier is then trained with the transformed training data. The query scRNA-Seq data will also be pair-transformed, and a classification score is generated for each query cell. (B) A representative example of the sixteen pairs of cross-platform scRNA-Seq training-query performance analyses. The Baron human pancreas scRNA-Seq dataset is the training and Murano human pancreas scRNA-Seq dataset is the query, were used to benchmark the performance of the five methods: SCN-TP, SCN-Base, SCMAP-cosine, SCMAP-pearson and SCMAP-spearman, using two quantitative metrics: mean AUPR and k. The training for each pair of the cross-platform comparisons was k-fold crossvalidated for ten times. Left: The barplot shows the mean and standard deviation of the classifier performance. Right: The classification scores are displayed with violin plot, where the x axis shows the true cell annotation, and the y axis label on the right-hand side shows the classifier category. (C) Mean and standard deviation of the classifier performance (Left: Kappa, right: mean AUPR) of cross-platform and cross-species scRNA-Seq classifiers.

After gene-pair selection, the training data is then transformed into a binary matrix and used to train a multi-class Random Forest classifier. Our training process also includes a step to generate, by random sampling of gene pair values, a set of transformed single cell profiles that are unlike any others in the training data. This ‘unknown’ category can be useful to identify query cells to which no class in the training data corresponds. Query scRNA-seq data undergoes the same TP transform. To measure the performance of the classifier throughout this study, we have used two assessment metrics: Cohen’s kappa (k), which measures agreement of categorical variables normalized for chance (Cohen, 1968), and mean area under the precision-recall curve (mean AUPR). As ground truth, we used a variety of gold standard data sets in which the cell identity is given as our base for comparison. To evaluate the performance of existing method SCMAP using mean AUPR, which requires a quantitative score, we used the intermediate outputs of Pearson, Spearman, and Cosine correlations.

### Performance of the SingleCellNet TP-RF classifier

We first set out to determine how the number of top pairs, the primary user-tunable parameter of our method, impacts classifier performance. In this and the other analyses in this section, we used the *Tabula Muris* 10x scRNA-Seq (TM-10x) dataset. First, we sampled 50 cells from each of the 32 defined cell types from this dataset for training top-pair SCN (SCN-TP) across a range of top pairs per cell type. Next, we used the 23,337 remaining cells as held-out validation data, finding that both k and mean AUPR plateaued when the number of top pairs was 10, corresponding to 320 total predictor genes (**Supp Fig. 1A**).

Next, we determined the effect of adjusting profiles based on stage of cell cycle, as this is sometimes considered an uninformative biological confounder of clustering analysis (Barron and Li, 2016). We evaluated performance in three scenarios: no cycle adjustment, adjustment of both training and validation data, and adjustment of validation data only, where adjustment is performed by regression on stage of cell cycle (Wolf et al., 2018). In addition to evaluating SCN-TP, we also trained and evaluated a version of SCN in which no top-pair transform is performed, but rather in which expression estimates of differentially expressed genes are used as predictors variables in training the Random Forest. We refer to this method as SCN-Base. In the scenario of no adjustments, SCN-TP and SCN-Base performed similarly (k 0.93 vs 0.94 and mean AUPR 0.95 vs 0.96). However, in the case of adjusting both the training and the validation data, the performance of SCN-TP held steady while that of SCN-Base plummeted (k 0.93 vs 0.29 and mean AUPR 0.94 vs 0.31). In the case in which only the validation data was adjusted for cell cycle, SCN-Base worsened further while SCN-TP remained stable (k 0.92 vs 0.05, mean AUPR 0.93 vs 0.19). This analysis suggests that SCN-TP is resilient to regressing on stage of cell cycle whereas other classifiers trained directly on expression levels are prone to performance degradation.

### Classifier performance across platforms and across species

There is wide diversity in scRNA-Seq methodologies, and the extent to which classifiers trained on data from one platform would be applicable to a query dataset from another, is unclear. We explored this by training classifiers and assessing their performance when applied to independent, well-annotated scRNA-Seq data from other studies of different scRNA-Seq platforms (**Supp Table 1**). We have assessed the classifier performance of all five methods with sixteen different pairs of cross-platform training and query data (**Supp Fig 1A**). As a representative analysis, we discuss here the results of using human pancreas cells profiled by inDrop as training data (Baron et al., 2016), and human pancreas cells profiled by CEL-Seq2 as the query (Muraro et al., 2016). SCN-TP had significantly higher mean AUPR compared to the SCMAP correlation methods and compared to SCN-Base (0.98 vs 0.60-0.88 vs 0.93, respectively) (**Fig 1B left**). SCN-TP and SCMAP had similar k, and both methods significantly outperformed SCN-Base (0.91 vs 0.89-0.93 vs 0.77). In typical scRNA-Seq studies, the data are clustered into groups of cells with similar profiles. If the clusters represent cell states or cell types that are robustly detectable across platforms and studies, then cells within the clusters should share high classification scores for the same category, and low scores for all other categories. To test this idea, we visualized the classification results as violin plots, where the major x-axis is the pancreas cell type as annotated in Murano *et al*, and the y-axis is the classification score of the indicated category (**Fig 1B, right**). Indeed, each cluster of the query data had one clear category with a maximal classification score. Interestingly, this analysis also illustrated that the SCN methods achieved a starker contrast in classification scores than SCMAP correlation methods (**Fig 1B right**), which may contribute to the lower mean AUPR of these methods. The performance results described above held true more generally. SCN methods had significantly higher mean AUPRs than SCMAP correlation methods in 16/16 analyses, whereas SCN had similar or higher k in 12/16 analyses (**Supp Fig 1A**).

Finally, we determined the performance of the methods when applied across species with five datasets: three of the pancreas and two of the central nervous system (**Supp Table 2**). To train these classifiers, we converted query dataset gene symbols to symbols of orthologs as determined by HCOP (Seal et al., 2011). Consistent with prior results, SCN-TP achieved significantly higher mean AUPR values, and either similar or higher k values than other methods (**Fig 1C**). The only exception was in one of the pancreas analyses in which SCN-Base achieved a similar mean AUPR and moderately higher k than SCN-TP.

Collectively, these analyses show that SCN-TP achieves a high, if not the highest, classifier performance across a range of conditions, including correction for cell cycle, differences in platforms, and differences in species.

### Example applications of SingleCellNet

Here we briefly describe one example application of SCN in perhaps the simplest use-case in which a user has performed a scRNA-Seq experiment and wants assign to the cells a putative identity. We used the Baron *et al* human pancreas data to train SCN-TP, and the Segerstolpe *et al* pancreas data as the query (**Fig 2A**). The primary visualization in SCN is a classification heatmap, in which the cells are represented as columns, the cell types of the classifier are represented as rows, and the multi-class classification scores are colored from black (low classification score) to yellow (high classification score). In this example, we ordered the query cells according to the annotation provided in the corresponding study (**Fig 2B top**). In addition to the fact that all of the previously annotated cells were appropriately classified, SCN also provided putative identities for the 43 pancreatic cells that had been previously “unclassified” in the original study: two gamma cells, nine alpha cells, three beta cells, three delta cells, one ductal cell, and two possible Schwann cells. The remaining cells have seemingly dual identities cells of either alpha and beta cells or alpha and gamma cells (**Fig 2B bottom**). In the online documentation for SCN and in the supplemental figures, we have included several other example applications, including a classification of fibroblasts reprogrammed to a neural-like identity (**Supp Fig 2**) (Tsunemoto et al., 2018), and a cross-species classification of peripheral blood mononuclear cells (**Supp Fig 3**) (Zheng et al., 2017). In these applications we have described additional ways to visualize the results, including violin classification plots, attribution plots, and dimensionality reduction based on classification results with UMAP.

**Figure 2.**
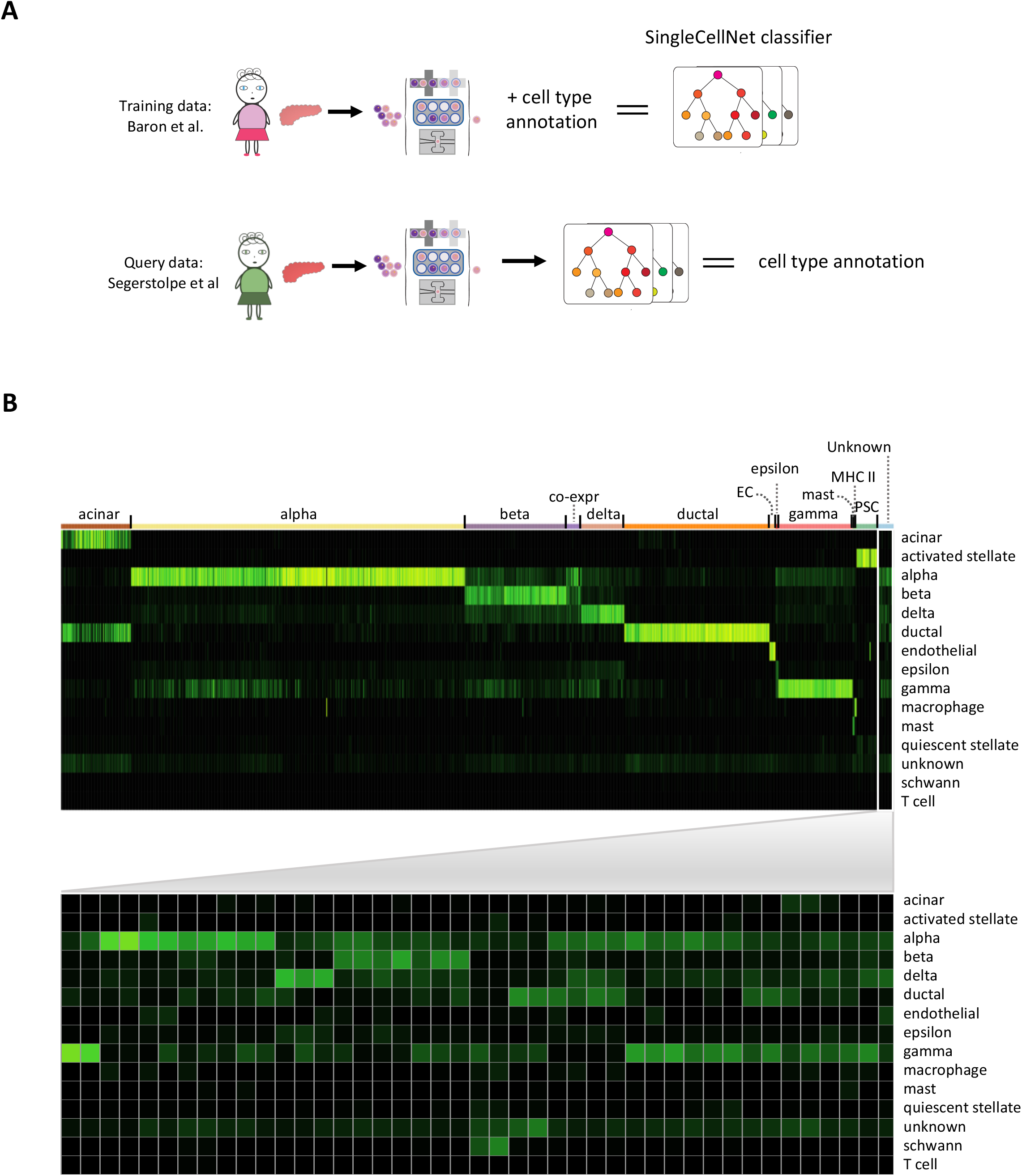
Application of SCN across scRNA-Seq platforms to determine the identity of unknown cells. (A) Baron adult human pancreatic scRNA-Seq data and its annotation provided by the authors are used to train SCN-TP classifier. Segerstolpe et al profiled adult human pancreatic scRNA-Seq data with a different scRNA-Seq technique was used as query data. (B) A classification heatmap is used to visualize the classification result of the Segerstolpe data on the Baron-based SCN-TP classifier. The annotated columns are labeled according the provided label from the Segerstolpe group. The row names indicate the cell-type specific classifier trained with the Baron data. The classification score for each column/cell sums up to 1, with a range from 0 (black) to 1 (yellow).

## Discussion

The demand for a robust method to quantify cell identity will grow as technologies to generate scRNA-Seq data proliferate and become more accessible. Here, we have shown that SCN quantifies cell identity in a manner that is robust to varying scRNA-Seq platforms and that is capable of classifying cells across species. To aid in the community’s use and improvement of SCN, we have made it available under an Open Source software license, and the code is accessible at Github: (http://github.com/pcahan1/singleCellNet/). Our documentation includes sections on training new classifiers, troubleshooting, expected computation time, and step-by-step procedures to reproduce the examples described here.

In contrast to other scRNA-Seq integration and comparison methods, we expect that SCN will be especially useful when a quantitative, rather than a binary, metric of identity is informative, or when the presence of shared cell types across datasets is unclear. One such application is the classification of engineered cell types (i.e. those derived through directed differentiation or direct conversion) in comparison to a reference dataset of *in* vivo-derived cells. As more data across developmental time points are accrued, we anticipate that SCN will provide a means to quantify not only the identity but also the stage of development and maturation of engineered cells.

### STAR Methods

Key Resources Table

| REAGENT or RESOURCE | Source | Identifier |
| --- | --- | --- |
| Tabula Muris scRNA-Seq data | (Tabula Muris Consortium et al., 2018) | https://figshare.com/projects/Tabula_Muris_Transcriptomic_characterization_of_20_organs_and_tissues_from_Mus_musculus_at_single_cell_resolution/27733 |
| Microwell-Seq scRNA-Seq data | (Han et al., 2018) | https://figshare.com/articles/MCA_DGE_Data/5435866 |
| Baron pancreas scRNA-Seq data | (Baron et al., 2016) | https://hemberg-lab.github.io/scRNA.seq.datasets/human/pancreas/ |
| Xin pancreas scRNA-Seq data | (Xin et al., 2016) | https://hemberg-lab.github.io/scRNA.seq.datasets/human/pancreas/ |
| Segerstolpe pancreas scRNA-Seq data | (Segerstolpe et al., 2016) | https://hemberg-lab.github.io/scRNA.seq.datasets/human/pancreas/ |
| Murano pancreas scRNA-Seq data | (Muraro et al., 2016) | https://hemberg-lab.github.io/scRNA.seq.datasets/human/pancreas/ |
| Zheng pbmc scRNA-Seq data | (Zheng et al., 2017) | https://support.10xgenomics.com/single-cell-gene-expression/datasets |
| Darminis brain scRNA-Seq data | (Darmanis et al., 2015) | https://www.ncbi.nlm.nih.gov/geo/query/acc.cgi?acc=GSE67835 |
| **Software and Algorithms** |  |  |
| R version 3.5.1 | R Foundation for Statistical Computing | https://www.r-project.org/ |
| SingleCellNet | This study | http://github.com/pcahan1/singleCellNet/ |

## Methods

### Building and assessing the classifier

Building a classifier begins with a preprocessed gene expression matrix and a pre-annotated metadata where each cell the gene expression matrix is annotated (**Supp Fig 4**). We demonstrate how to use the SCN pipeline with the example provided in our online README (http://github.com/pcahan1/singleCellNet/), where the Tabula Muris 10X dataset was used as training and the Park *et al* dataset was used as query (Park et al., 2018). First, we found the intersection of genes between the training data and the query data prior to training the SCN-TP classifier. Then, we randomly selected 100 cells (ncell = 100) per cell type from the entire training data to train the SCN-TP classifier and reserved the remaining cells to measure the classifier’s performance (**Step 1**). The subsetted training data was then down-sampled to 1500 counts per cell (total = 1.5e3), scaled up such that the total expression per cell was 10000 (xFact=1e4), and log-transformed. Based on the annotation (dLevel = “newAnn”), we found the top ten (topX = 10) most differentially expressed genes per cell type (**Step 2a**), then we ranked top ten gene-pair per cell type (topX = 10) from those genes (**Step 2b**). To optimize memory usage, we have parallelized the ranking process, where we examined sets of gene pairs in chunks of 5000 (sliceSize=5000) (**Step 2b**). The preprocessed training data was then transformed according to the selected gene pairs (**Step 3**), and was used to build a multi-class SCN-TP classifier of 1000 trees (ntrees = 1000) (**Step 4**). Additionally, we created 100 randomized cell expression profiles (nrand = 100) to train up a “rand” category in the SCN-TP classifier, which can help in cases where some cell types that are present in the query data are not included in the training data (**Step 2b**). After the SCN-TP classifier was built, we transformed the remaining held-out data according to the top gene pairs selected (**Step 5a**), along with another 100 randomized cells (numRand = 100). To assess the performance of the classifier, we applied it to the transformed held-out data (**Step 5b**) using Precision-Recall curves, k, and mean AUPR (**Step 5c-e**). This is a crucial quality control step as it will indicate the optimal performance that can be expected from the classifier. If the classifier performs poorly on held out data, then the user should troubleshoot the training procedure beginning with the scRNA-Seq data annotation.

### Classifying query data

Once we determined that the classifier performed well, then we applied to the transformed external query data (Park et al., 2018) with top-pairs selected from the optimized training data (**Step 6**), and classified it with the SCN-TP classifier (**Step 7**). We displayed the results by i) classification heatmap, ii) UMAP, iii) attribution plot, and iv) classification violin plot (**Step 8**). To facilitate the visualization of the classification result, we created a named vector with each cell’s true identity (sla), and appended to it the randomized cells (slaRand), which we had created earlier. The attribution plot was generated with classification output (ct_score = classRes_val_all), metadata for the held-out data (stTest), number of random cells created when querying (nRand = 100), the column in metadata that stored the true identity of the held-out cells (dLevel = “newAnn”), and sample/cell names (sid = “cell”). Similar to the input for generating the attribution plot, we visualized the classification result with UMAP using the top 5 principal components based on the classification results (topPC = 5).

### Notes to users

The quality and annotation of the training data are critical to building reliable classifiers. We recommend to start training a classifier with 10-20 distinct cell types, and to iteratively add more cell types and assess classifier performance. Obviously, the user should not attempt to assess a classifier with the query data as the true identity of the query data is unknown.

## Author Contributions

YT and PC designed the computational pipeline and wrote the manuscript. PC conceived the and supervised the project.

## Acknowledgments

This work was supported by the National Institutes of Health under grant R35GM124725 to PC the Biochemistry, Cellular, and Molecular Biology Program T32 Training grant T32GM007445 to YT. We would also like to thank members of the Cahan lab members, especially Emily Su, Emily Lo, Dan Peng, Ray Cheng, and Abby Spangler, for providing feedback and support.

## Supplementary Information

**Supplementary Figure 1.**
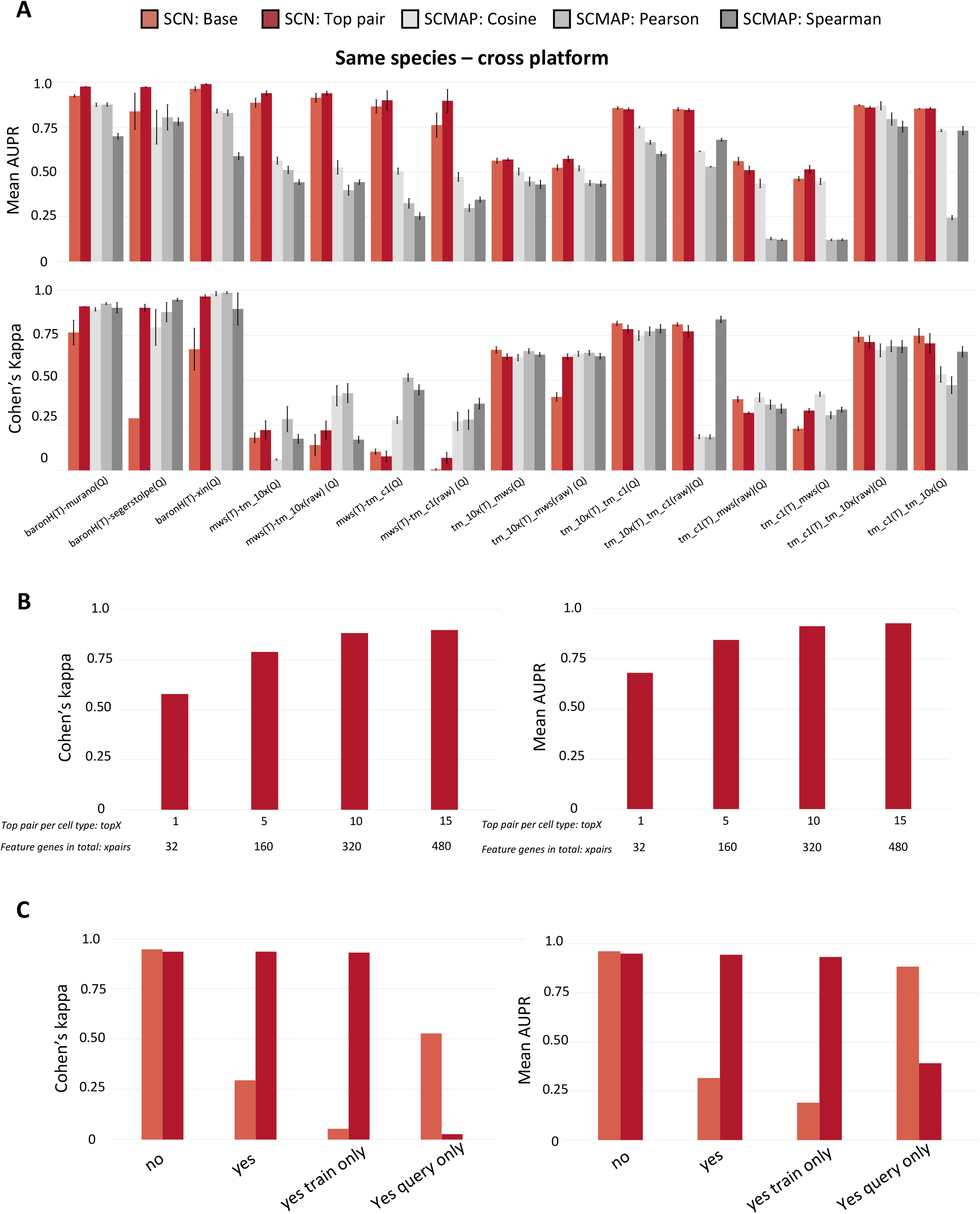
SCN parameter and performance correlation. (A) Sixteen pairs of cross-platform scRNA-Seq training-query datasets were used to benchmark the performance of the five methods: SCN-TP, SCN-Base, SCMAP-cosine, SCMAP-pearson and SCMAP-spearman, using two quantitative metrics: mean AUPR and k. The training for each pair of the cross-platform comparisons was bootstrapped ten times. The barplot exhibits the classifier performance of all ten training by displaying the average AUPR and standard deviation as error bar. (B) The Tabula Muris 10x dataset cross-validation was used to test how the number of top-pairs used in SCN can influence its performance. We have tested 1 top pair per cell type (32 genes in total), 5 top pairs per cell type (160 genes in total), 10 top pairs per cell type (320 genes in total), and 15 top pair per cell type (480 genes in total). The performance of SCN plateaus at 10 top-pairs per cell type for both mean AUPR and k. (C) How cell cycle regression may affect SCN performance. We have examined four conditions where 1) both training and query data have not been regressed for cell cycle, 2) both training and query data have regressed out cell cycle effect regressed out, 3) only training data has cell cycle effect regressed out. 4) only query data has cell cycle effect regressed out.

**Supplementary Figure 2.**
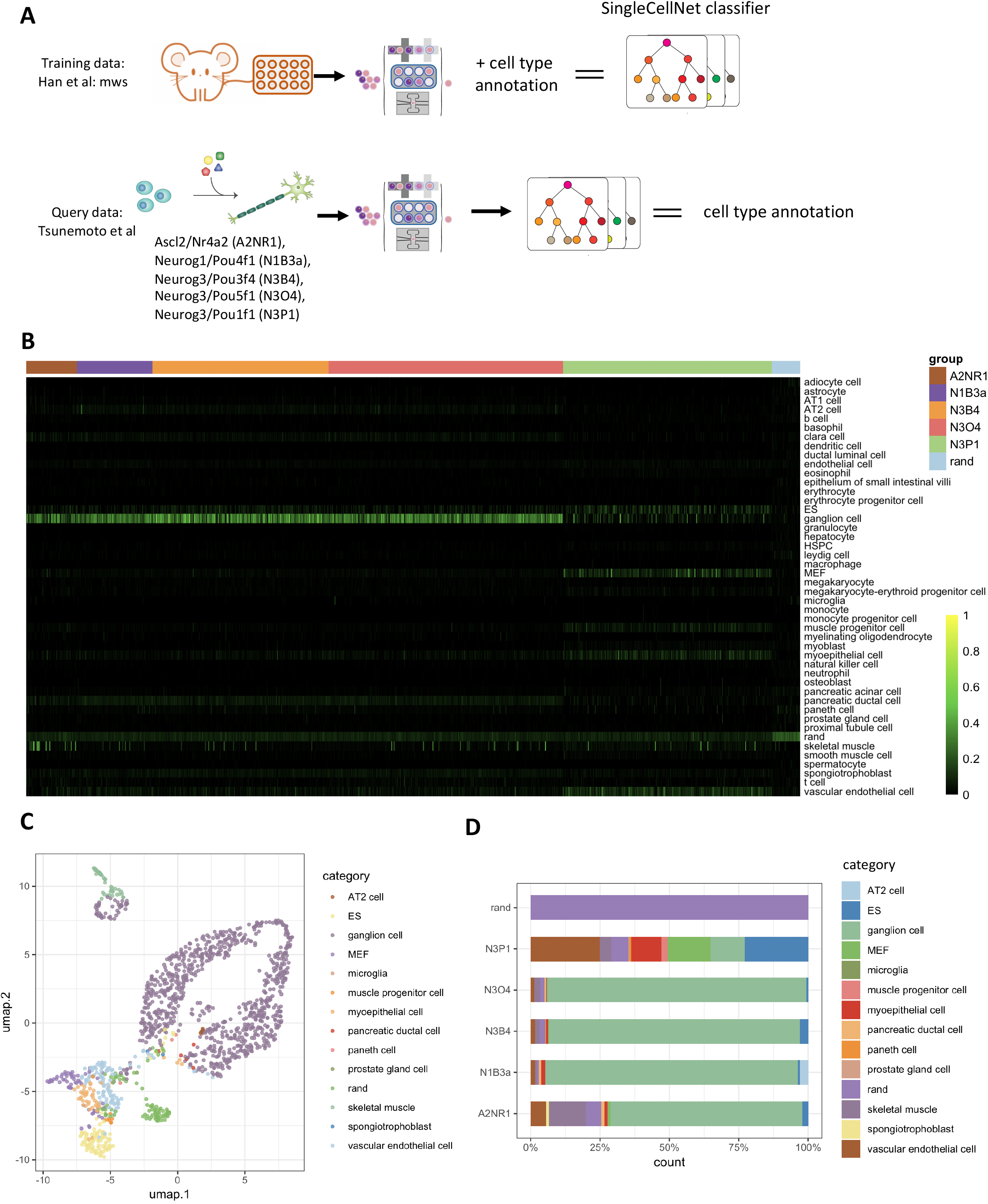
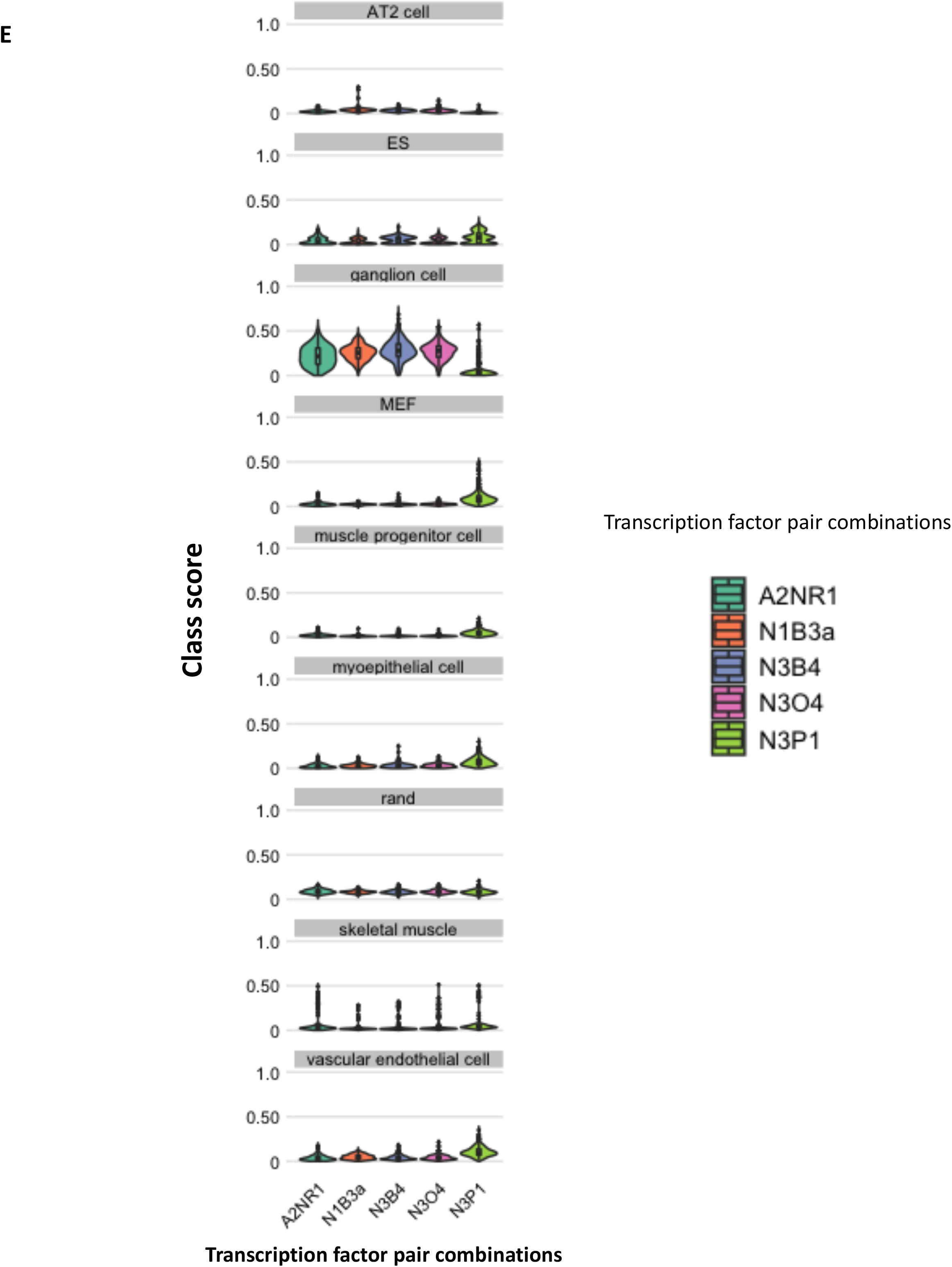
Application of SCN in quantitative assessment of direct reprogramming protocols. (A) We used a subset of adult Microwell-Seq scRNA-Seq data and its annotations to train a cell-type specific classifier. The Tsunemoto’s screening experiment, profiling scRNA-Seq of induced neurons (iNs) by application of transcription factor combinations, is used as query data. (B) The columns annotate the five different transcription factor pairs used to generate each iN profile, namely *Neurog3/Pou5f1* (N3O4), *Neurog3/Pou3f4* (N3B4), *Neurog1/Pou4f1* (N1B3a), *Ascl2/Nr4a2(A1NR1)* and *Nuerog3/Pou1f1* (N3P1). (C) The classification score can also be visualized with UMAP, where each cell is colored by its category or classification result of the highest score. (D) To obtain a more comprehensive understanding of the composition of cells in each reprogramming experiment, an attribution plot can be used with the row showing the experimental conditions and the column denoting the percentage count. Each cell is colored by its category or classification result of the highest score. (E) The classification score is visualized with violin plot, where x-axis displays the transcription factor pair combinations and y-axis is the range of classification score in that combination, and the plot is faceted by the classifier categories.

**Supplementary Figure 3.**
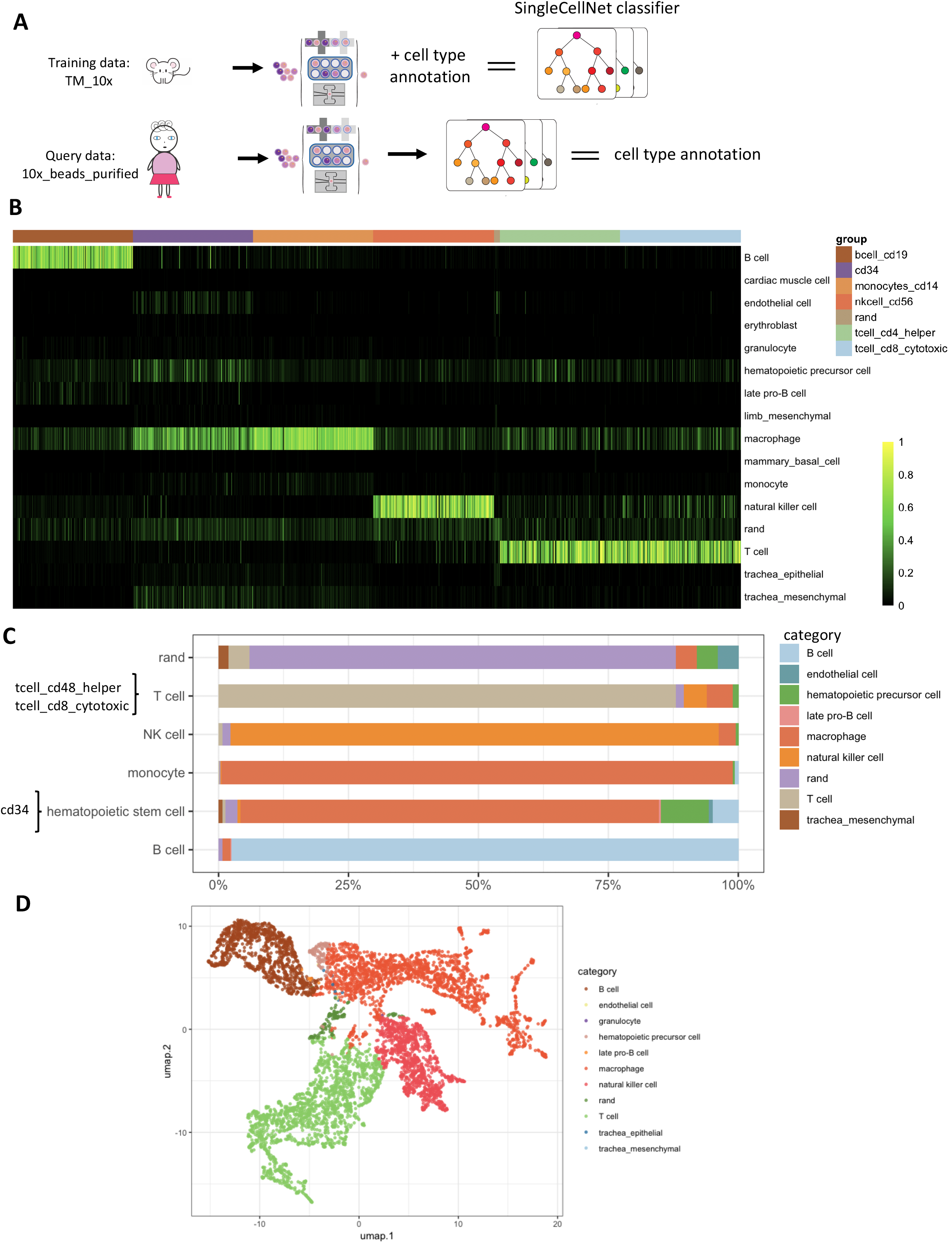
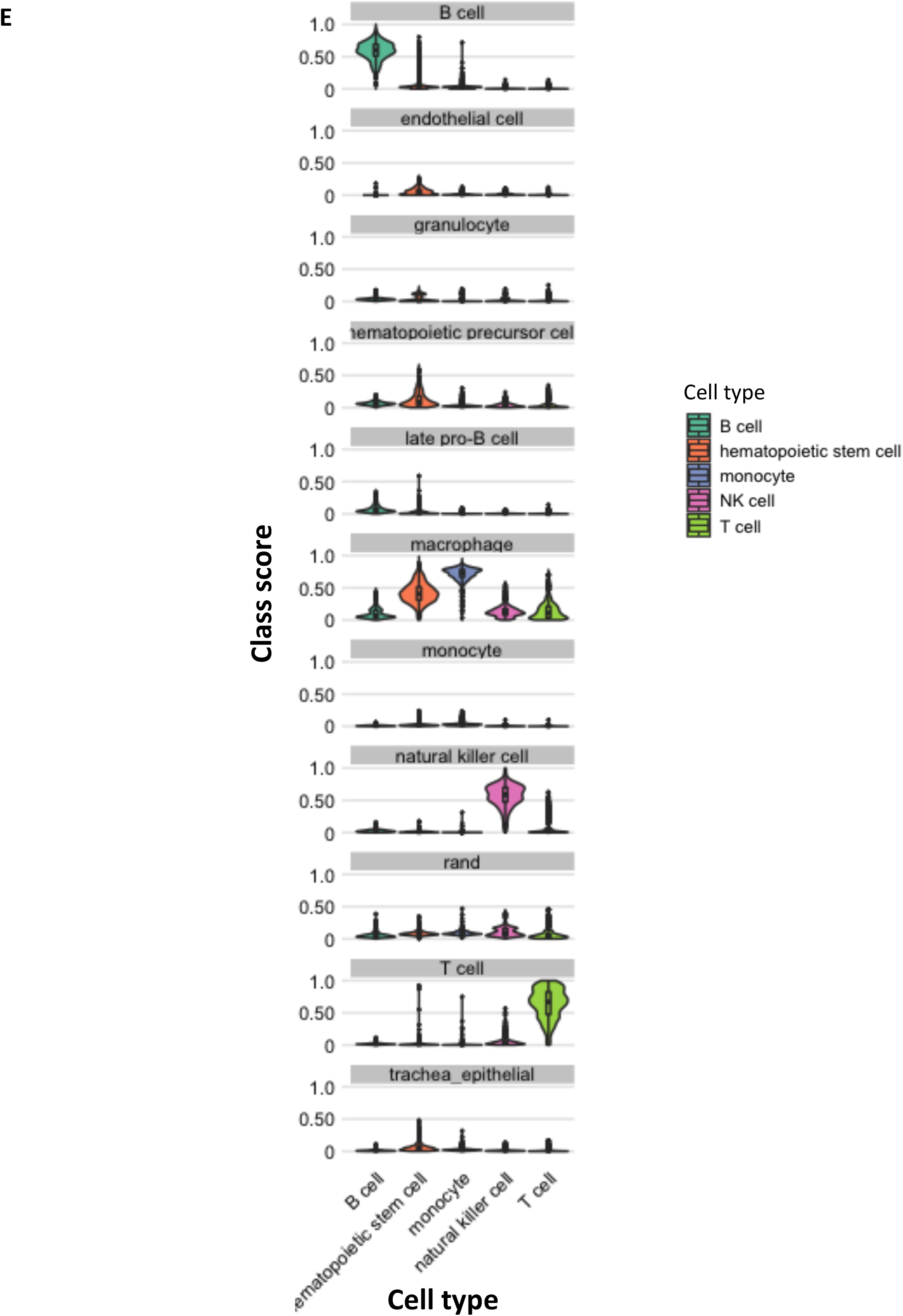
Application of SCN in cross-platform and cross-species analysis (A) The hematopoietic-lineage related subset of the Tabula Muris 10x scRNA-Seq data and its annotations are used as training data. The bead-purified 10x sequencing of the human hematopoietic lineage cells are used as query. (B) The classification heatmap shows that human B cells, monocytes, nature killer cells, CD4 T cells and CD8 T cells are well-classified by the mouse TP-RF classifier. (C) This cross-platform and cross-species analysis can also be summarized with an attribution plot, showing that despite some cross-classification signals due to the similarity, most human hematopoietic cells are accurately classified. (D) A side-by-side comparison of UMAP plots displaying the SCN-determined classification of the highest score versus the original annotation from bead-purified methods. (E) The classification score is visualized with violin plot, where x-axis shows the annotation of the query cells provided in the original studies, and y-axis is the range of classification score in that combination, and the plot is then faceted by the classifier categories.

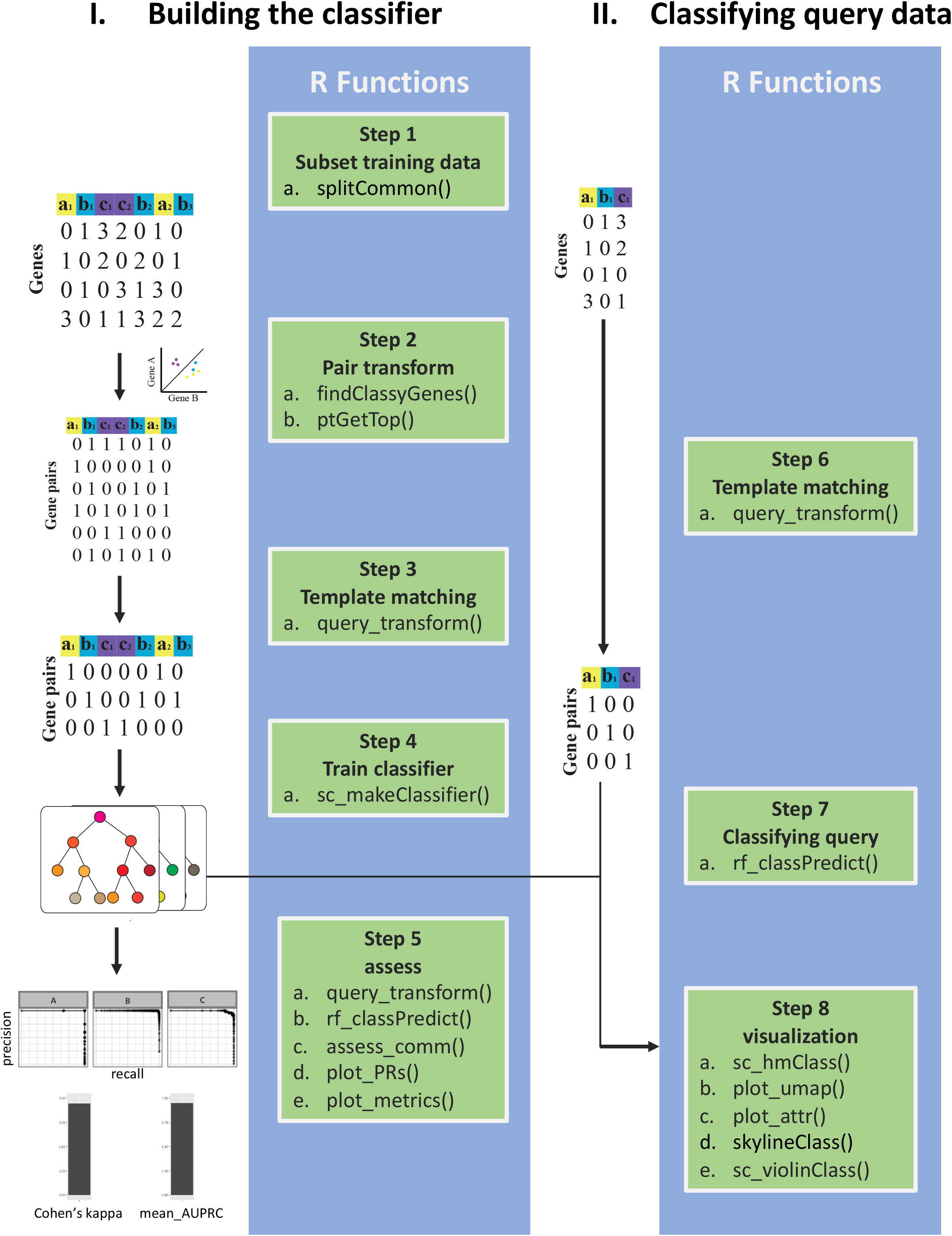

**Supplementary Table 1:** scRNA-Seq training and query datasets used for cross-platform comparison

| Cross-platform<br>same species<br>analysis | Training Data | Query Data |
| --- | --- | --- |
| 1 | **baron**<br>8596 human pancreatic<br>cells, inDrop | **murano**, 2120 adult human pancreatic<br>cells, CEL-Seq2 |
| 2 |  | **segerstolpe**, 2209 adult human<br>pancreatic cells, Smart-Seq2 |
| 3 |  | **xin**, 1492 human pancreatic cells, C1 |
| 6 - 7 | **tm_10x**,<br>1599 adult mouse cells<br>across 32 cell types, 10x | **mws \| mws (raw)**, 6477 mouse cells<br>across 125 cell types, Microwell-seq |
| 8 - 9 |  | **tm_c1 \| tm_c1 (raw)** tabula muris facs<br>data, 40620 adult mouse cells across 69<br>cell types, c1 |
| 10 - 11 | **tm_c1**,<br>3182 adult mouse cells<br>across 69 cell types, c1 | **tm_10x \| tm_10x (raw)** (tabula muris<br>10x data), 24936 adult mouse cells<br>across 32 cell types, 10x |
| 11 - 12 |  | **mws \| mws (raw)**, 6477 adult mouse<br>cells across 125 cell types, Microwell-Seq |
| 13 - 14 | **mws**,<br>6477 adult mouse cells<br>across 125 cell types,<br>Microwell-seq | **tm_10x \| tm_10x (raw)** tabula muris<br>10x data, 24936 adult mouse cells<br>across 32 cell types, 10x |
| 15 – 16 |  | **tm_c1 \| tm_c1 (raw)** tabula muris facs<br>data, 40620 adult mouse cells across 69<br>cell types, c1 |

\*(raw) indicates the query expression matrix was used in raw count form

**Supplementary Table 2:** scRNA-Seq training and query datasets used for cross-species comparison

| Cross-platform cross-species analysis | Training Data | Query Data |
| --- | --- | --- |
| 1 | **baron**<br>1886 adult mouse pancreatic cells, inDrop | **murano**, 2120 adult human pancreatic cells, CEL-Seq2 |
| 2 |  | **segerstolpe**, 2209 adult human pancreatic cells, Smart-Seq2 |
| 3 |  | **xin**, 1492 human pancreatic cells, C1 |
| 4 - 5 | **mws**<br>4038 adult mouse brain cells, Microwell-Seq | **darminis \| darminis (raw)**, 331 adult human brain cells, C1 |

\*(raw) indicates the query expression matrix was used in raw count form

